# Limited correspondence in visual representation between the human brain and convolutional neural networks

**DOI:** 10.1101/2020.03.12.989376

**Authors:** Yaoda Xu, Maryam Vaziri-Pashkam

## Abstract

Convolutional neural networks (CNNs) have achieved very high object categorization performance recently. It has increasingly become a common practice in human fMRI research to regard CNNs as working model of the human visual system. Here we reevaluate this approach by comparing fMRI responses from the human brain in three experiments with those from 14 different CNNs. Our visual stimuli included original and filtered versions of real-world object images and images of artificial objects. Replicating previous findings, we found a brain-CNN correspondence in a number of CNNs with lower and higher levels of visual representations in the human brain better resembling those of lower and higher CNN layers, respectively. Moreover, the lower layers of some CNNs could fully capture the representational structure of human early visual areas for both the original and filtered real-world object images. Despite these successes, no CNN examined could fully capture the representational structure of higher human visual processing areas. They also failed to capture that of artificial object images in all levels of visual processing. The latter is particularly troublesome, as decades of vision research has demonstrated that the same algorithms used in the processing of natural images would support the processing of artificial visual stimuli in the primate brain. Similar results were obtained when a CNN was trained with stylized object images that emphasized shape representation. CNNs likely represent visual information in fundamentally different ways from the human brain. Current CNNs thus may not serve as sound working models of the human visual system.

**Significance Statement:** Recent CNNs have achieved very high object categorization performance, with some even exceeding human performance. It has become common practice in recent neuroscience research to regard CNNs as working models of the human visual system. Here we evaluate this approach by comparing fMRI responses from the human brain with those from 14 different CNNs. Despite CNNs’ ability to successfully perform visual object categorization like the human visual system, they appear to represent visual information in fundamentally different ways from the human brain. Current CNNs thus may not serve as sound working models of the human visual system. Given the current dominating trend of incorporating CNN modeling in visual neuroscience research, our results question the validity of such an approach.

## INTRODUCTION

Recent hierarchical convolutional neural networks (CNNs) have achieved amazing human-like object categorization performance (Kriegeskorte, 2015; Yamins & Dicarlo, 2016; Rajalingham, et al., 2018; Serre, 2019). Additional research revealed that representations formed in early and late layers of the network track those of the human early and later visual processing regions, respectively (Khaligh-Razavi & Kriegeskorte, 2014; Güçlü & van Gerven, 2015; Cichy et al., 2016; Eickenberg et al., 2017). Together with results from monkey neurophysiological studies, CNNs have been regarded by some as the current best models of the primate visual system (e.g., Cichy & Kaiser, 2019; Kubilius et al., 2019).

Despite these excitements, researchers debated on what should be considered as a valid model of the human visual system. With its large number of parameters that do not closely follow the known architecture of the primate visual brain, some argue that CNNs may not be able to serve as models of the brain (Kay, 2018; Serre, 2019). But perhaps CNNs are not meant to capture the details of the neural processing, but rather, the computations that transform visual information from image pixels to recognizable objects at Marr’s algorithmic level (Marr, 1981). Even in this regard, due to a lack of detailed understanding of information processing in different CNN layers, CNNs appear more like black boxes than well understood information processing systems (Serre, 2019). Recent research effort has revealed that the best CNNs could only explain about 50% of the response variance of macaque V4 and IT (Cadieu et al., 2014; Yamins et al., 2014; Kar et al. 2019; Bashivan et al., 2019). The correlation between CNNs and brain/behavior has been shown to be fairly low in several cases (Kheradpisheh et al., 2016; Karimi-Rouzbahani et al., 2017; Rajalingham, et al., 2018). Others have reported that the kind of features used in object recognition differ between the brain and CNNs (Ballester & de Araujo, 2016, Ulman et al., 2016; Gatys et al., 2017; Baker et al., 2018; Geirhos et al., 2019). Adversarial images are particularly troublesome for CNNs as they minimally impact human object recognition performance but significantly decrease CNN performance (Serre, 2019).

Even though differences are noted between CNNs and the primate brain, it has recently become common practice in human fMRI studies to compare fMRI measures to CNN outputs (e.g, Long et al., 2018; Bracci et al., 2019; King et al., 2019). This is mainly based on the key fMRI finding showing that representations formed in early and late layers of the CNN could track those of the human early and later visual processing regions, respectively (Khaligh-Razavi & Kriegeskorte, 2014; Güçlü & van Gerven, 2015; Cichy et al., 2016; Eickenberg et al., 2017). This was established in one approach using linear transformation to link individual fMRI voxels to the units of CNN layers through training and cross-validation (Güçlü & van Gerven, 2015; Eickenberg et al., 2017). While this is a valid approach, it is computationally costly and requires large amounts of training data to map the large number of fMRI voxels to the even larger number of CNN units. Others have bypassed this direct voxel-to-unit mapping, and instead examined the correspondence in visual representational structures between the human brain and CNNs using representational similarity analysis (RSA, Kriegeskorte & Kievit, 2013). With this approach, both Khaligh-Razavi and Kriegeskorte (2014) and Cichy et al. (2016) reported a close correspondence in representational structure between early and late human visual areas and early and later CNN layers, repectively. Khaligh-Razavi and Kriegeskorte (2014) additionally showed that such correlations exceeded the noise ceiling for both brain regions, indicating that the representations formed in a CNN could fully capture those of human visual areas.

These human findings are somewhat at odds with results from neurophysiological studies showing that the current best CNNs can only capture about 50% of the response variance of macaque V4 and IT (Cadieu et al., 2014; Yamins et al., 2014; Kar et al. 2019). Additionally, Khaligh-Razavi and Kriegeskorte (2014) and Cichy et al. (2016) were underpowered by using an event-related fMRI design and anatomically defined visual areas, raising concerns regarding the robustness of their findings. Most importantly, none of the above fMRI studies tested the representation of object images beyond the unaltered real-world object images. As human participants have no trouble recognizing filtered real-world object images (such as the high spatial frequency (SF) components of an image, which resemble a line drawing), it would be critical to know how well a close brain-CNN correspondence could be generalized to these filtered object images. Decades of visual neuroscience research has successfully utilized simple and artificial visual stimuli to uncover the complexity of visual processing in the primate brain, showing that the same algorithms used in processing natural images would manifest themselves in the processing of artificial visual stimuli. If CNNs are to be used as working models of the primate visual brain, it is equally critical to test whether a close brain-CNN correspondence exists for the processing of artificial objects.

Given that recently published fMRI studies and many more ongoing fMRI projects rely on the brain-CNN correspondence being robust and generalizable across image sets, here we reevaluated this finding with several different image sets using an fMRI block design in functionally defined human visual areas. Because the RSA approach is simple and elegant, allowing easy comparisons of multiple fMRI data sets with multiple CNNs, and because a noise ceiling can be easily derived to quantify the degree of brain-CNN correspondence, we will use this approach in the present study. By comparing the visual representational structure from three fMRI experiments and 14 different CNNs, we show that although visual processing in lower and higher human visual areas indeed resemble those of lower and higher CNN layers, respectively, only lower CNN layers could fully capture the visual representational structure of real-world objects in lower human visual areas. CNNs, as a whole, fully capture neither the representations of real-world objects at higher levels of visual processing nor those of artificial objects at either level of processing. The same results were obtained regardless of whether a CNN was trained with real-world object images or stylized object images that emphasized shape representation. These results, together with those from other recent studies, point to fundamental differences between the human brain and CNNs and put significant limitations on how CNNs may be used to model visual processing in the human brain.

## RESULTS

In this study, we reexamined previous finding that showed a close brain-CNN correspondence in visual processing (Khaligh-Razavi & Kriegeskorte, 2014; Güçlü & van Gerven, 2015; Cichy et al., 2016; Eickenberg et al., 2017). We noticed that the two studies that used the RSA approach were underpowered in two aspects. First, both Khaligh-Razavi and Kriegeskorte (2014) and Cichy et al. (2016) used an event-related fMRI design, known to produce low SNR. This can be seen in the low brain and CNN correlation values reported, with the highest correlation being less than .2 in both studies. While Cichy et al. (2016) did not calculate the noise ceiling, thus making it difficult to assess how good the correlations were, the lower bounds of the noise ceiling were around .15 to .2 in Khaligh-Razavi and Kriegeskorte (2014), which was fairly low. Second, both studies defined human brain regions anatomically rather than functionally on each individual participant. This could affect the reliability of fMRI responses, potentially contributing to the low noise ceiling and low correlation obtained. Here we took advantage of several existing fMRI data sets that overcome these drawbacks and compared visual processing in the human brain with 14 different CNNs. These data sets were collected while human participants viewed both unfiltered and filtered real-world object images, as well as artificial object images. This allowed us to test not only the robustness of brain-CNN correlation, but also its generalization across different image sets.

Our fMRI data were collected with a block design in which responses were averaged over a whole block of multiple exemplars to increase SNR. In three fMRI experiments, human participants viewed blocks of sequentially presented cut-out images on a grey background and pressed a response button whenever the same image repeated back to back (Figure 1a). Each image block contained different exemplars from the same object category. A total of eight real-world natural and manmade object categories were used, including bodies, cars, cats, chairs, elephants, faces, houses, and scissors (Vaziri-Pashkam & Xu, 2019; Vaziri-Pashkam et al., 2019). In Experiment 1, both the original images and controlled version of the same images were shown (Figure 1b). Controlled images were generated using the SHINE technique to achieve spectrum, histogram, and intensity normalization and equalization across images from the different categories (Willenbockel et al., 2010). In Experiment 2, the original, high and low SF content of an image from six of the eight real-world object categories were shown (Figure 1b). In Experiment 3, both the images from the eight real-world image categories and images from nine artificial object categories (Op de Beeck et al., 2008) were shown (Figure 1b).

**Figure 1.**
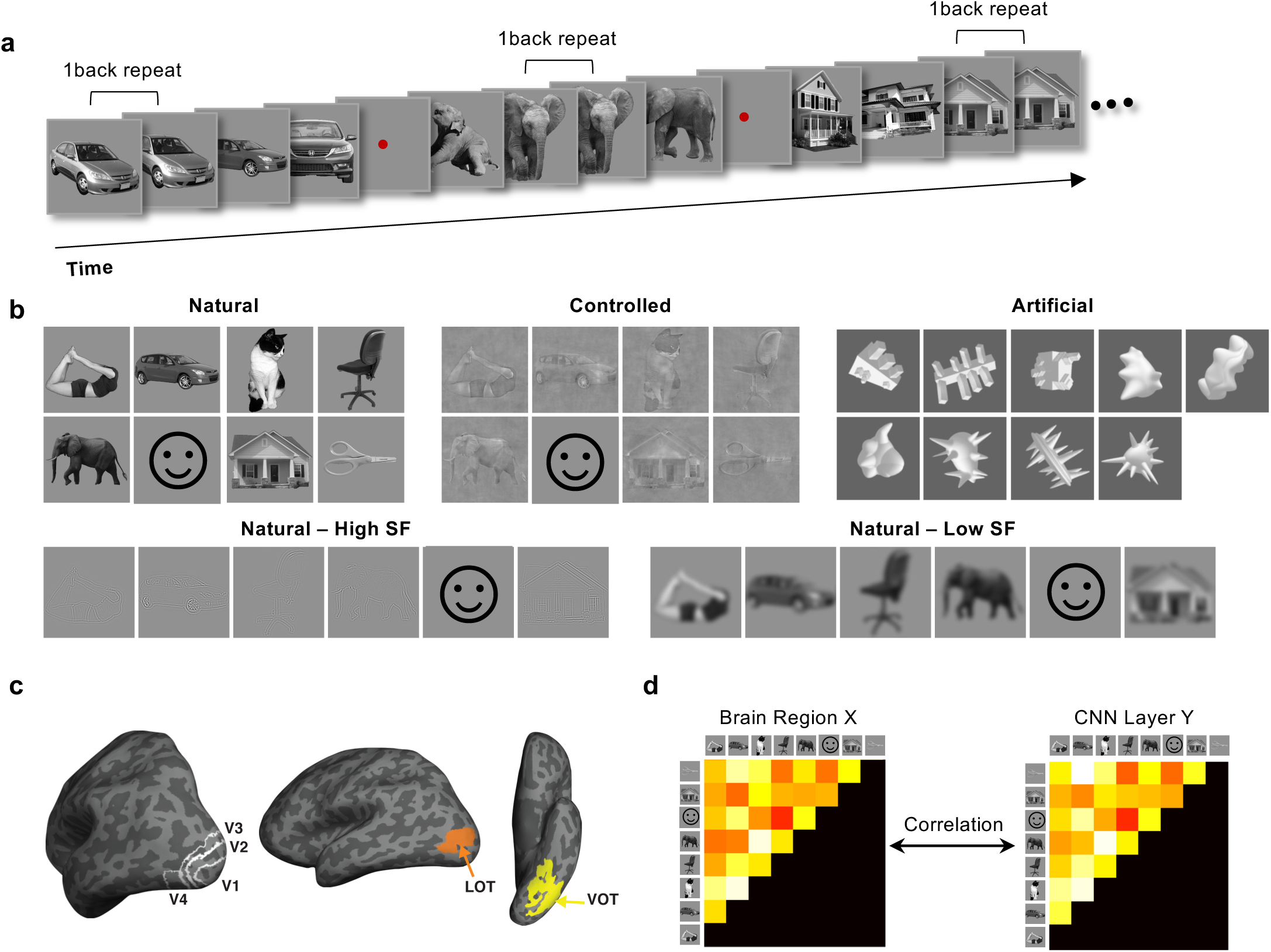
**(A)** An illustration of the block design paradigm used. Participants performed a one-back repetition detection task on the images. An actual block in the experiment contained 10 images with two repetitions per block. See Methods for more details. **(B)** The stimuli used in the three fMRI experiments. Experiment 1 included the original and the controlled images from eight real-world object categories. Experiment 2 included the images from six of the eight real-world object categories shown in the original, high SF, and low SF format. Experiment 3 included images from the same eight real-world object categories and images from nine artificial object categories. Each category contained ten different exemplars varying in identity, pose, and viewing angle to minimize the low-level image similarities among them. Due to copyright issues, an example of the actual face images used in the experiments is not shown here but is depicted by a cartoon face for illustration purposes. **(C)** The brain regions examined. They included topographically defined early visual areas V1 to V4 and functionally defined higher object processing regions LOT and VOT. **(D)** The representational similarity analysis used to compare the representational structural between the brain and CNNs. In this approach, a representation dissimilarity matrix was first formed by computing all pairwise Euclidean distances of fMRI response patterns or CNN layer output for all the object categories. The off-diagonal elements of this matrix were then used to form a representational dissimilarity vector. Lastly, these dissimilarity vectors were correlated between each brain region and each sampled CNN layer to assess the similarity between the two.

For a given brain region, we averaged fMRI responses from a block of exemplars of the same category and extracted the beta weights (from a general linear modal) for the entire block from each voxel as the fMRI response pattern for that object category. Following this, fMRI response patterns were extracted for each category from six independently defined visual regions along the human OTC. They included early visual areas V1 to V4 and higher visual object processing regions in lateral occipito-temporal (LOT) and ventral occipito-temporal (VOT) cortex (Figure 1c). LOT and VOT have been considered as the homologue of the macaque inferio-temproal (IT) cortex involved in visual object processing (Orban et al., 2004). Their responses have been shown to correlate with successful visual object detection and identification (Grill-Spector et al. 2000; Williams et al., 2007) and their lesions have been linked to visual object agnosia (Goodale et al.,1991; Farah, 2004).

The 14 CNNs we examined here included both shallower networks, such as Alexnet, VGG16 and VGG 19, and deeper networks, such as Googlenet, Inception-v3, Resnet-50 and Resnet-101 (Table 1). We also included a recurrent network, Cornet-S, that has been shown to capture the recurrent processing in macaque IT cortex with a shallower structure (Kubilius et al., 2019; Kar et al., 2019). This CNN has been recently argued to be the current best model of the primate ventral visual processing regions (Kar et al., 2019). All CNNs were pretrained with ImageNet images (Deng et al., 2009). To understand how the specific training images would impact CNN representations, besides CNNs trained with ImageNet images, we also examined Resnet-50 trained with stylized ImageNet images (Geirhos et al., 2019). Following a previous study (O’Connor et al., 2018), we sampled from 6 to 11 mostly pooling layers of each CNN (see Table 1 for the specific CNN layers sampled). Pooling layers were selected because they typically mark the end of processing for a block of layers before information is pooled and passed on to the next block of layers. We extracted the response from each sampled CNN layer for each exemplar of a category and then averaged the responses from the different exemplars to generate a category response, similar to how an fMRI category response was extracted. Following Khaligh-Razavi and Kriegeskorte (2014) and Cichy et al. (2014), using RSA, we compared the representational structure of real-world and artificial object categories between the different CNN layers to those obtained in the human visual processing regions.

**Table 1.**
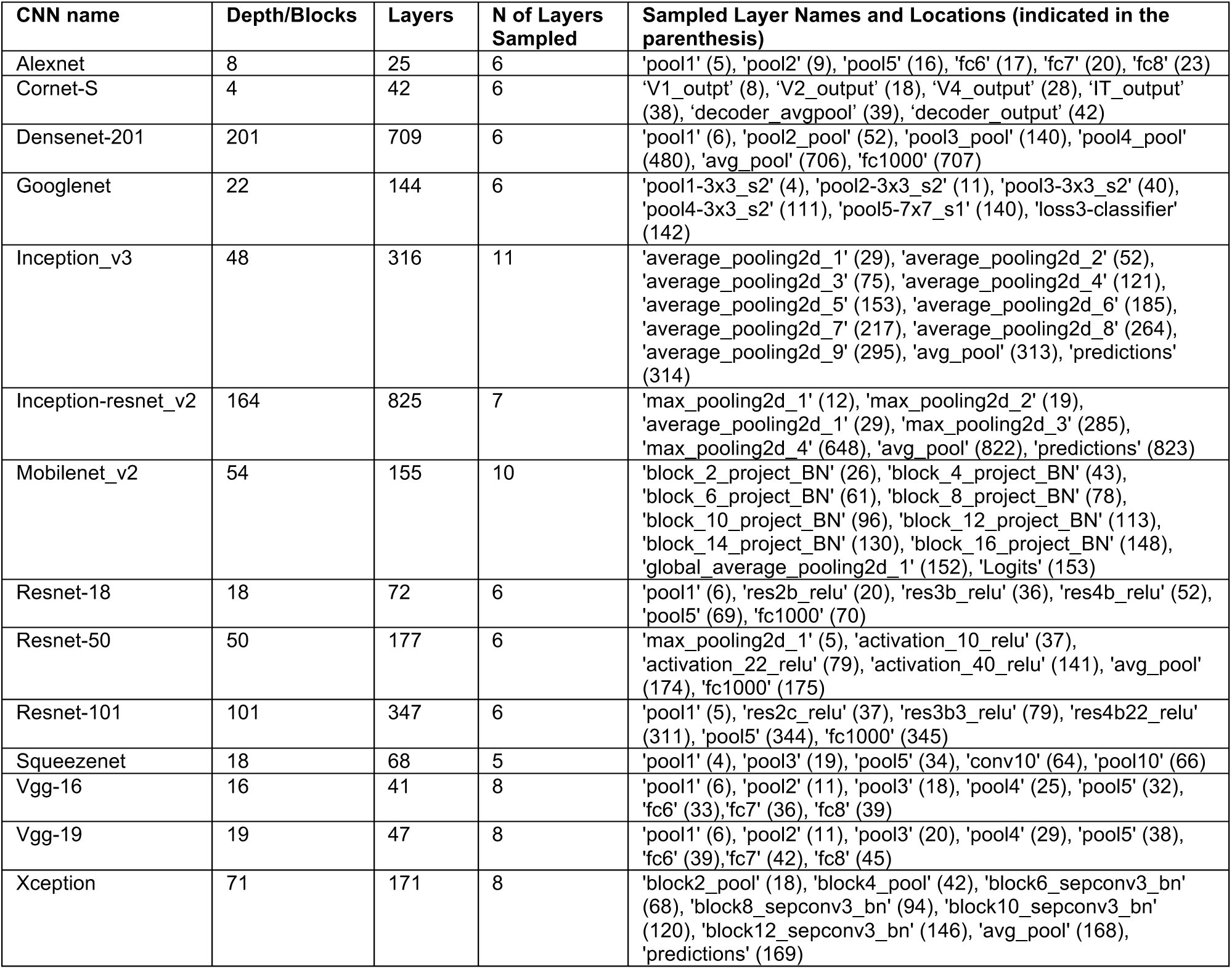
The CNNs and the layers examined in this study.

| CNN name | Depth/Blocks | Layers | N of Layers Sampled | Sampled Layer Names and Locations (indicated in the parenthesis) |
| --- | --- | --- | --- | --- |
| Alexnet | 8 | 25 | 6 | 'pool1' (5), 'pool2' (9), 'pool5' (16), 'fc6' (17), 'fc7' (20), 'fc8' (23) |
| Cornet-S | 4 | 42 | 6 | 'V1_output' (8), 'V2_output' (18), 'V4_output' (28), 'IT_output' (38), 'decoder_avgpool' (39), 'decoder_output' (42) |
| Densenet-201 | 201 | 709 | 6 | 'pool1' (6), 'pool2_pool' (52), 'pool3_pool' (140), 'pool4_pool' (480), 'avg_pool' (706), 'fc1000' (707) |
| Googlenet | 22 | 144 | 6 | 'pool1-3x3_s2' (4), 'pool2-3x3_s2' (11), 'pool3-3x3_s2' (40), 'pool4-3x3_s2' (111), 'pool5-7x7_s1' (140), 'loss3-classifier' (142) |
| Inception_v3 | 48 | 316 | 11 | 'average_pooling2d_1' (29), 'average_pooling2d_2' (52), 'average_pooling2d_3' (75), 'average_pooling2d_4' (121), 'average_pooling2d_5' (153), 'average_pooling2d_6' (185), 'average_pooling2d_7' (217), 'average_pooling2d_8' (264), 'average_pooling2d_9' (295), 'avg_pool' (313), 'predictions' (314) |
| Inception-resnet_v2 | 164 | 825 | 7 | 'max_pooling2d_1' (12), 'max_pooling2d_2' (19), 'average_pooling2d_1' (29), 'max_pooling2d_3' (285), 'max_pooling2d_4' (648), 'avg_pool' (822), 'predictions' (823) |
| Mobilenet_v2 | 54 | 155 | 10 | 'block_2_project_BN' (26), 'block_4_project_BN' (43), 'block_6_project_BN' (61), 'block_8_project_BN' (78), 'block_10_project_BN' (96), 'block_12_project_BN' (113), 'block_14_project_BN' (130), 'block_16_project_BN' (148), 'global_average_pooling2d_1' (152), 'Logits' (153) |
| Resnet-18 | 18 | 72 | 6 | 'pool1' (6), 'res2b_relu' (20), 'res3b_relu' (36), 'res4b_relu' (52), 'pool5' (69), 'fc1000' (70) |
| Resnet-50 | 50 | 177 | 6 | 'max_pooling2d_1' (5), 'activation_10_relu' (37), 'activation_22_relu' (79), 'activation_40_relu' (141), 'avg_pool' (174), 'fc1000' (175) |
| Resnet-101 | 101 | 347 | 6 | 'pool1' (5), 'res2c_relu' (37), 'res3b3_relu' (79), 'res4b22_relu' (311), 'pool5' (344), 'fc1000' (345) |
| Squeezenet | 18 | 68 | 5 | 'pool1' (4), 'pool3' (19), 'pool5' (34), 'conv10' (64), 'pool10' (66) |
| Vgg-16 | 16 | 41 | 8 | 'pool1' (6), 'pool2' (11), 'pool3' (18), 'pool4' (25), 'pool5' (32), 'fc6' (33), 'fc7' (36), 'fc8' (39) |
| Vgg-19 | 19 | 47 | 8 | 'pool1' (6), 'pool2' (11), 'pool3' (20), 'pool4' (29), 'pool5' (38), 'fc6' (39), 'fc7' (42), 'fc8' (45) |
| Xception | 71 | 171 | 8 | 'block2_pool' (18), 'block4_pool' (42), 'block6_sepconv3_bn' (68), 'block8_sepconv3_bn' (94), 'block10_sepconv3_bn' (120), 'block12_sepconv3_bn' (146), 'avg_pool' (168), 'predictions' (169) |

### The existence of brain-CNN correspondence for representing real-world object images

In Experiments 1 and 2, we had two goals: first, To replicate and quantify the previous reported brain-CNN correspondence for representing real-world object images; and second, to test whether this finding can be generalized to filtered real-world images. As human participants had no trouble recognizing our filtered real-world object images, if there is indeed a close brain-CNN correspondence, such a correspondence should be preserved for the representation of the filtered versions of the real-world object images.

To compare the representational structure between the human brain and CNNs, in each brain region examined, we first calculated pairwise Euclidean distances of the z-normalized fMRI response patterns for the object categories in each experiment, with shorter Euclidean distance indicating greater similarity between a pair of fMRI response patterns. From these pairwise Euclidean distances, we constructed a category representational dissimilarity matrix (RDM, see Figure 1d) for each of the six brain regions examined. Likewise, from the z-normalized category responses of each sampled CNN layer, we calculated pairwise Euclidean distances among the different categories to form a CNN category RDM for that layer. We then correlated category RDMs between brain regions and CNN layers using Spearman rank correlation following Nili et al. (2014) and Cichy et al. (2016) (Figure 1d). A Spearman rank correlation compares the representational geometry between the brain and a CNN without requiring the two having a strict linear relationship. All our results remained the same when Spearman correlation was applied and when correlation measures, instead of Euclidean distance measures, were used (see Supplemental Figures 1 to 4).

**Figure 4.**
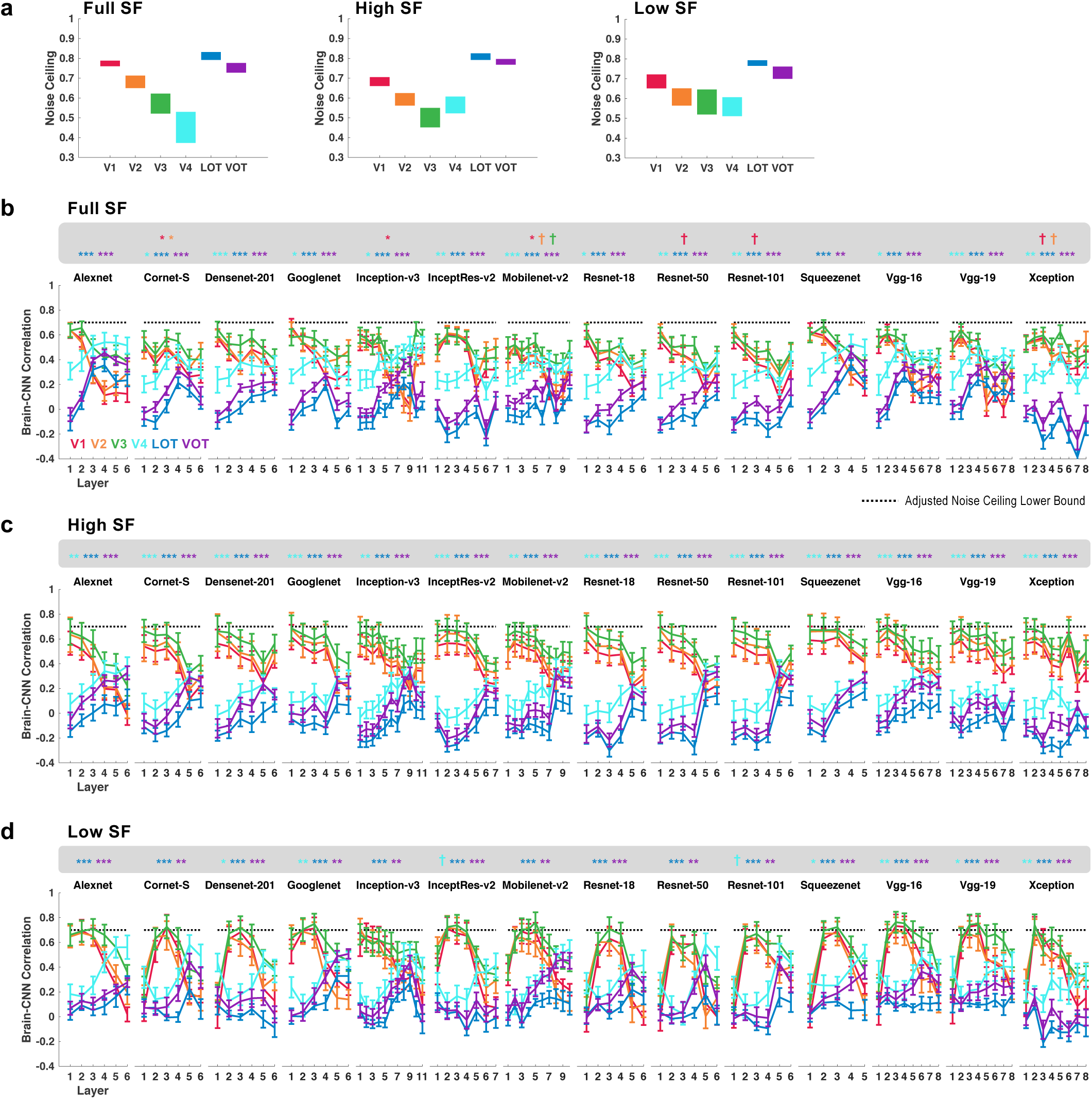
Quantifying the brain-CNN correspondence in Experiment 2 with original, high SF and low SF images from real-world object categories. (A) The upper and lower bounds of noise ceiling of the fMRI responses for each image condition. (B) to (D) RDM correlations of each brain region with each sampled layer in each CNN for the Original, high SF and low SF images, respectively. The lower bounds of the noise ceiling from all brain regions were set to 0.7. † p < .1, * *p < .05*, ** *p < .01*, *\*\*\* p < .001*.

Previous studies have reported a correspondence in representation between lower and higher CNN layers to lower and higher visual processing regions, respectively (Khaligh-Razavi & Kriegeskorte, 2014; Cichy et al., 2016). To evaluate the presence of such a correspondence in our data, for each CNN examined, we identified in each human participant, the CNN layer that showed the best RDM correlation with each of the six brain regions included. We then assessed whether the resulting layer numbers increased from low to high visual regions using Spearman rank correlation. If a close CNN-brain correspondence in representation exists, then the Fisher-transformed correlation coefficient of this Spearman rank correlation should be significantly above zero at the group level (all stats reported were corrected for multiple comparisons for the number of comparisons included in each experiment using the Benjamini–Hochberg procedure at false discovery rate q = .05, see Benjamini & Hochberg, 1995).

In Experiment 1, we contrasted original real-world object images with the controlled version of these images. Figure 2a (purple lines) shows the averaged CNN layer corresponding to each brain region in each CNN when original real-world objects images were shown (the significance levels of the CNN-brain correspondence were marked with asterisks at the top of each plot). Here 10 out of the 14 CNNs examined showed a significant CNN-brain correspondence. The same correspondence was also seen when controlled images were shown in Experiment 1, with 11 of the 14 CNNs showing a significant CNN-brain correspondence (Figure 2a blue lines).

**Figure 2.**
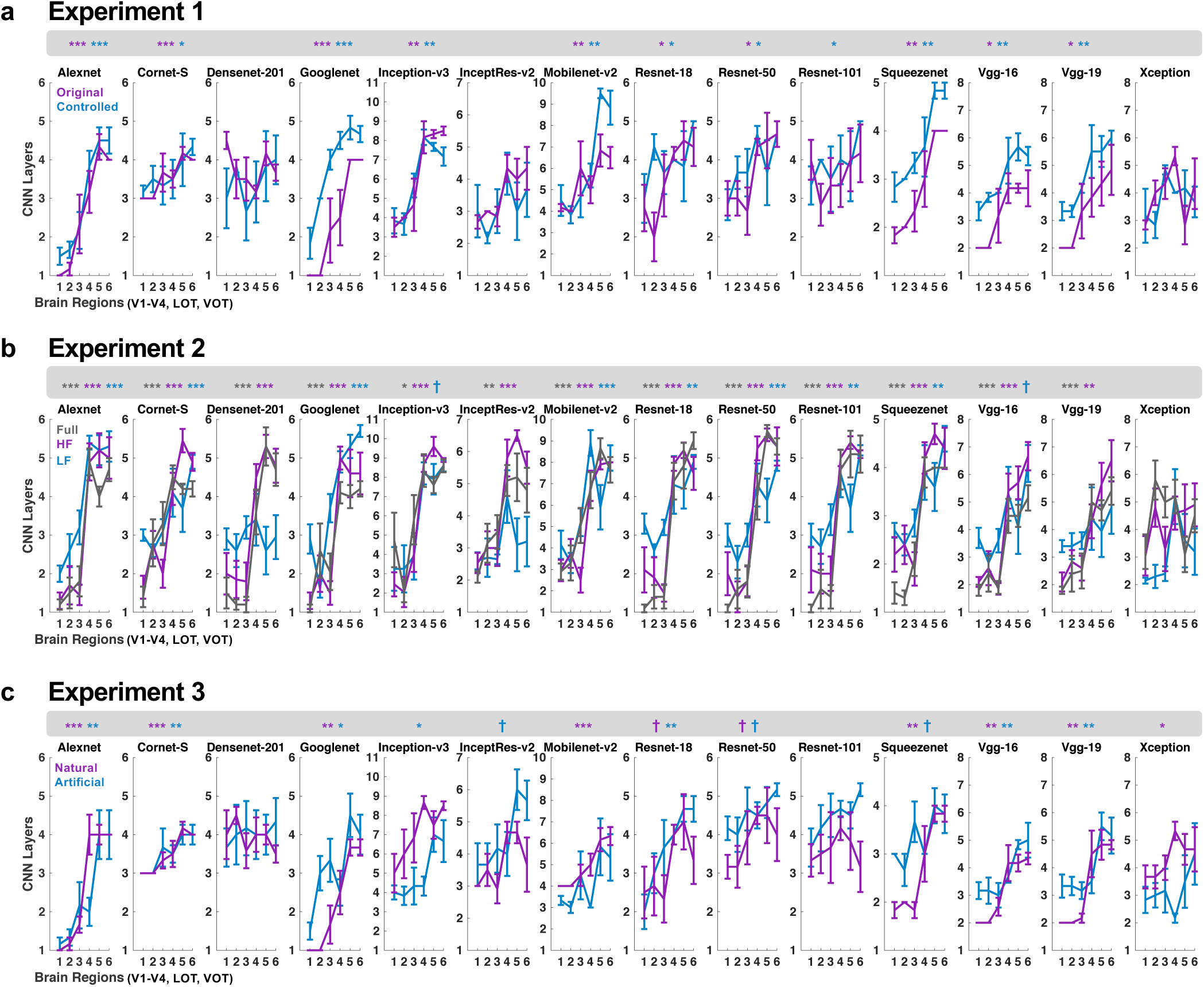
Evaluating the presence of the brain-CNN correspondence in their representational structures for **(A)** Experiment 1, in which original and controlled images from the real-world categories were shown, **(B)** Experiment 2, in which original, high and low SF components of the images from the real-world categories were shown, and **(C)** Experiment 3, in which the unaltered images from both the real-world and artificial object categories were shown. To evaluate the brain-CNN correspondence, we identified in each human participant, the CNN layer that showed the best RDM correlation with each of the six brain regions. We then assessed whether the resulting layer numbers linearly increased from low to high visual regions using Spearman rank correlation. Finally, we tested the significance of the resulting correlation coefficients (Fisher-transformed) at the group level. All statistical tests were corrected for multiple comparisons for the number of image conditions included in an experiment using the Benjamini–Hochberg procedure. * *p < .05*, ** *p < .01*, *\*\*\* p < .001*.

In Experiment 2, we contrasted original real-world images with the high and low SF component version of these images. For the original images, we replicated the findings from Experiment 1, with 13 out of the 14 CNNs showing a significant brain-CNN correspondence (Figure 2b, grey lines). The same correspondence was also present in 13 CNNs for the high SF images, and in 8 CNNs for the low SF images (Figure 2b, purples and blue lines). In fact, Alexnet, Cornet-S, Googlenet, Mobilenet-v2, Resnet-18, Resnet-50 and Squeezenet showed a significant brain-CNN correspondence for all five image sets across Experiments 1 and 2.

These results replicated previous findings using the RSA approach (Khaligh-Razavi & Kriegeskorte, 2014; Cichy et al., 2016) and showed that there indeed existed a linear correspondence between CNN and brain representations, such that representations in lower and higher visual areas better resemble those of lower and higher CNN layers, respectively. Importantly, such a brain-CNN correspondence was generalizable to filtered real-world object images.

### Quantifying the amount of brain-CNN correspondence for representing real-world object images

A linear correspondence between CNN and brain representations, however, only tells us that lower CNN layers are relatively more similar to lower than higher visual areas and that the reverse is true for higher CNN layers. It says nothing about the amount of similarity. To assess this, we evaluated how successful the category RDM from a CNN layer could capture the RDMs from a brain region. To do so, we first obtained the reliability of the category RDM in a brain region across the group of human participants by calculating the lower and upper bounds of the fMRI noise ceiling (Nili et al., 2014). Overall, the lower bound of our fMRI noise ceilings were much higher across the different ROIs in the two experiments than those of Khaligh-Razavi and Kriegeskorte (2014) (Figures 3a and 4a). This indicated that the object category representational structures were fairly similar and consistent among the different participants.

**Figure 3.**
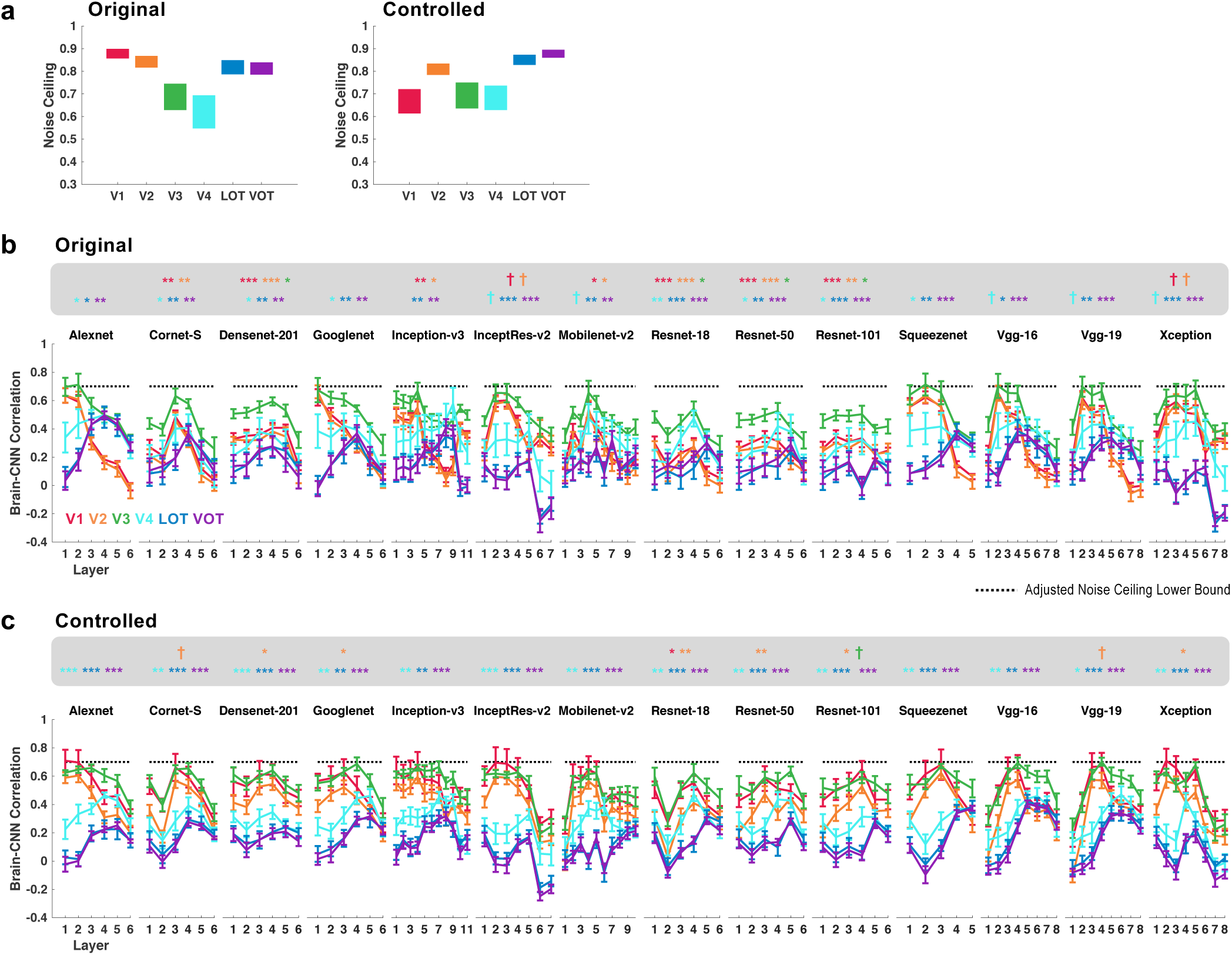
Quantifying the brain-CNN correspondence in Experiment 1 with original and controlled images from real-world object categories. (A) The upper and lower bounds of noise ceiling of the fMRI responses for each image condition. (B) and (C) RDM correlations of each brain region with each sampled layer in each CNN for the Original and Controlled images, respectively. Because the lower bound of the noise ceiling varied somewhat among the different brain regions, for illustration purposes, while maintaining the differences between the CNN and brain correlations with respect to their lower bound noise ceilings, the lower bounds of the noise ceiling from all brain regions were set to 0.7. † p < .1, * *p < .05*, ** *p < .01*, *\*\*\* p < .001*.

If the category RDM from a CNN layer successfully captures that from a brain region, then the correlation between the two should exceed the lower bound of the fMRI noise ceiling. Because the lower bound of the fMRI noise ceiling varied somewhat among the different brain regions, for illustration purposes, in all the data plots shown (Figures 3 to 6), while the differences between the CNN and brain correlations with respect to the noise ceiling lower bounds were maintained, the noise ceiling lower bounds from all brain regions were set to 0.7. For the original real-world object images in Experiment 1, the correlation between lower visual areas and CNN layers reached the fMRI noise ceiling in five CNNs (Figure 3b), including Alexnet, Googlenet, Squeenzenet, Vgg16, and Vgg19 (Fisher-transformed distance to fMRI noise ceiling lower bound, *ps* > .1; see the asterisks marking the significance level for the highest correlation between each brain region and a CNN layer at the top of each plot; all p values reported were corrected for multiple comparisons for the 6 brain regions included using the Benjamini–Hochberg procedure at false discovery rate q = .05, see Benjamini & Hochberg, 1995). However, no correlation between higher visual areas LOT and VOT in any CNN layer reached the noise ceiling (Fisher-transformed distance to noise ceilings, *p*s < .05, corrected). The same pattern of results was seen when the controlled images were used in Experiment 1 (Figure 3c), with several CNNs able to fully capture the representational structure of visual processing in lower visual areas but none able to do so for higher visual areas. We obtained similar results for the original, high SF and low SF images in Experiment 2 (Figures 4b to 4d). Here again, a number of CNNs were able to fully capture the RDM of lower visual areas, but none could do so for higher visual areas. All these results remained the same when correlations, instead of Euclidean distance measures were used. Additionally, these results held when Pearson, instead of Spearman correlations were applied (see Supplemental Figures 1 to 4), indicating that early layers of some of the CNNs were able to fully capture the object representational structure in early visual areas in a linear manner.

Together, these results showed that, across the two experiments, although lower layers of a number of CNNs could fully capture lower level visual processing of both the original and filtered real-world object images in the human brain, none could do so for higher level neural processing of these images.

### The brain-CNN correspondence for representing artificial object images

Previous comparisons of brain and CNN visual processing have focused entirely on the representation of real-world objects. Decades of visual neuroscience research, however, has successfully utilized simple and artificial visual stimuli to uncover the complexity of visual processing in the primate brain, showing that the same algorithms used in processing natural images would manifest themselves in the processing of artificial visual stimuli. If CNNs are to be used as working models of the primate visual brain, it would be critical to test if this principle applies to CNNs.

In Experiment 3, we contrasted the processing of both real-world object images and artificial object images. As in Experiments 1 and 2, the processing of real-world object images showed a consistent CNN-brain correspondence in 8 out of the 14 CNNs tested (Figure 2c, purple lines). Interestingly, the same correspondence was also obtained in 8 CNNs when artificial objects were shown, with lower visual representations better resembling those of lower than higher CNN layers and the reverse being true for higher visual representations (Figures 2c). In fact, across Experiments 1 to 3, Alexnet, Cornet-S and Googlenet were the three CNNs showing a consistent brain-CNN correspondence across all our image sets including original and filtered real-world object images, as well as artificial object images.

Nevertheless, when we compared RDM correlation between the brain and CNNs, we again found that, for real-world object images, while some of the CNNs were able to fully capture the visual representational structure of lower level visual processing in the human brain, no CNNs could do so for higher level visual processing in the brain (Figure 5b). For artificial object images, while there still existed a significant difference in RDM correlation between higher brain regions and higher CNN layers in a majority of the CNNs, the close RDM correlation between lower visual areas and lower CNN layers was no longer present in any of the 14 CNNs examined (Figure 5c). In other words, as a whole, CNNs performed much worse in capturing visual processing of artificial than real-world object images in the human brain.

**Figure 5.**
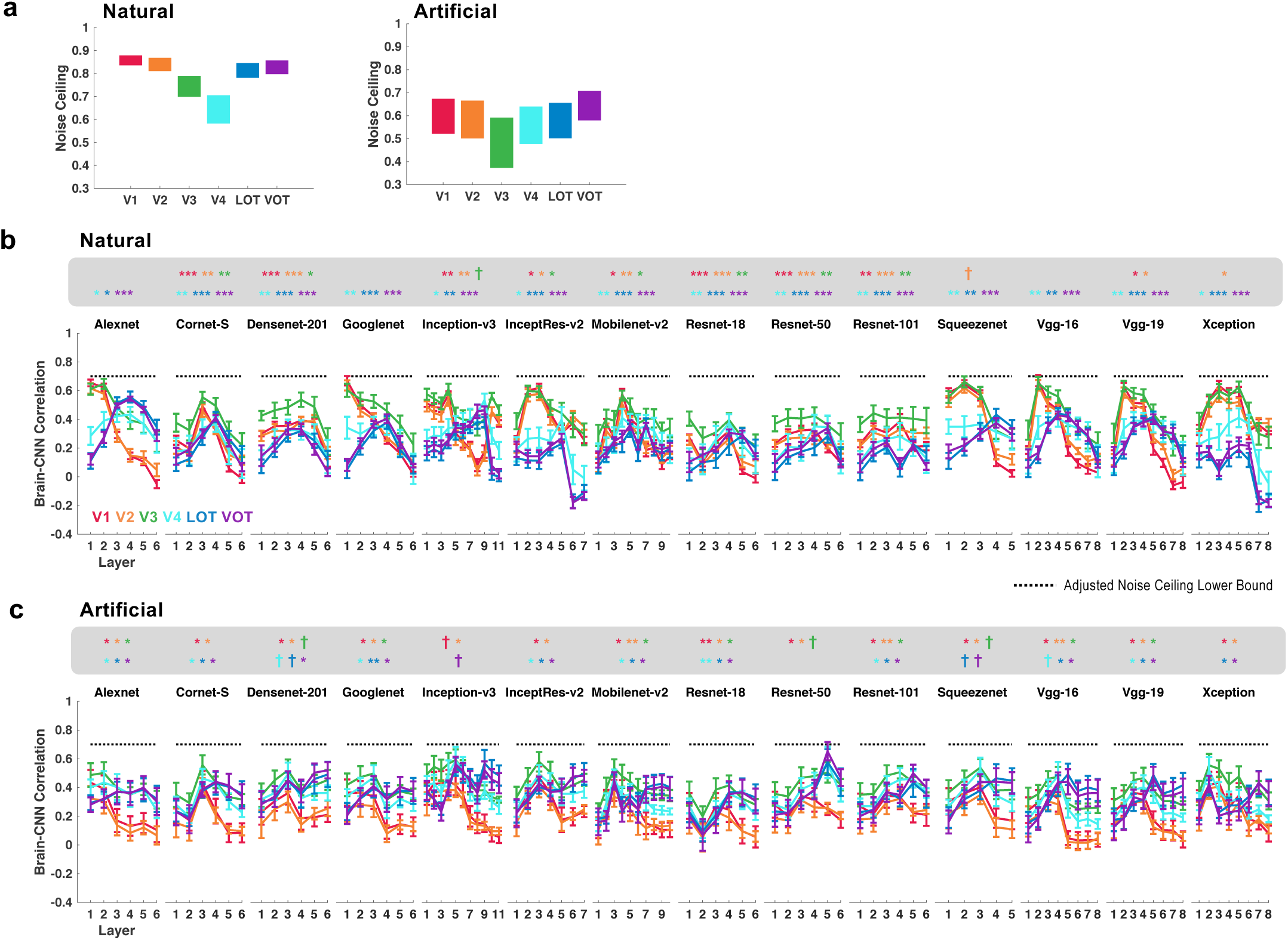
Quantifying the brain-CNN correspondence in Experiment 3 with images from real-world and artificial object categories. (A) The upper and lower bounds of noise ceiling of the fMRI responses for each image condition. (B) and (C) RDM correlations of each brain region with each sampled layer in each CNN for the real-world and artificial object images, respectively. The lower bounds of the noise ceiling from all brain regions were set to 0.7. † p < .1, * *p < .05*, ** *p < .01*, *\*\*\* p < .001*.

Overall, despite the presence of a linear correspondence in representation between the brain and some CNNs, taking both the linear correspondence and RDM correlation into account, none of the CNNs examined here could fully capture lower and higher levels of visual processing for artificial objects. This is particularly troublesome given that a number of CNNs were able to fully capture lower level visual processing of real-world objects in the human brain. Given that the same algorithms used in the processing of natural images have been shown to support the processing of artificial visual stimuli in the primate brain, CNNs’ inability to do so limit them as sound working models that can help us understand the nature of visual processing in the human brain.

### The effect of training a CNN on original vs stylized image-net images

Although CNNs are believed to explicitly represent object shapes in the higher layers (Kriegeskorte, 2015; LeCun et al., 2015; Kubilius et al., 2016), emerging evidence suggests that CNNs may largely use local texture patches to achieve successful object classification (Ballester & de Araujo, 2016, Gatys et al., 2017) or local rather than global shape contours for object recognition (Baker et al., 2018). In a recent demonstration, CNNs were found to be poor at classifying objects defined by silhouettes and edges, and when texture and shape cues were in conflict, classifying objects according to texture rather than shape cues (Geirhos et al., 2019; see also Baker et al., 2018). However, when Resnet-50 was trained with stylized ImageNet images in which the original texture of every single image was replaced with the style of a randomly chosen painting, object classification performance significantly improved, relied more on shape than texture cues, and became more robust to noise and image distortions (Geirhos et al., 2019). It thus appears that a suitable training data set may overcome the texture bias in standard CNNs and allow them to utilize more shape cues.

Here we tested if the category RDM in a CNN may become more brain-like when a CNN was trained with stylized ImageNet images. To do so, we examined the representations formed in Resnet-50 pretrained with three different procedures (Geirhos et al., 2019): trained only with the stylized ImageNet Images (RN50-SIN), trained with both the original and the stylized ImageNet Images (RN50-SININ), and trained with both sets of images and then fine-tuned with the stylized ImageNet images (RN50-SININ-IN). For comparison, we also included Resnet-50 trained with the original ImageNet images (RN50-IN) that we tested before.

Despite minor differences, the category RDM correlations between brain regions and CNN layers were remarkably similar whether or not Resnet-50 was trained with the original or the stylized ImageNet images: all were still substantially different from those of the human visual regions (Supplemental Figure 5). If anything, RN50-IN showed a better brain-CNN correspondence in a few cases than the other training conditions. Thus, despite the improvement in object classification performance with the inclusion of the stylized images, how Resnet-50 represented object categories and encodes visual features did not appear to change. The incorporation of stylized ImageNet images likely forced Resnet-50 to use long-range structures rather than local texture structures, but without fundamentally changing how images were computed. The inability of Resnet-50 to exhibit brain-like category RDM was therefore unlikely due to the inadequacy of the training data used to train the network, but rather some fundamental differences between the two systems.

## DISCUSSION

Recent CNNs have achieved very high object categorization performance, with some even exceeding human performance (Serre, 2019). It has become common practice in recent human fMRI research to regard CNNs as a working model of the human visual system. In particular, previous fMRI studies have reported that representations formed in earlier and later layers of the CNN could track those of the human earlier and later visual processing regions in fMRI studies, respectively (Khaligh-Razavi & Kriegeskorte, 2014; Güçlü & van Gerven, 2015; Cichy et al., 2016; Eickenberg et al., 2017). This finding has inspired a host of subsequent fMRI studies to use CNNs as models of the human visual system and compare fMRI measures with CNN layer outputs (e.g, Bracci et al., 2019; King et al., 2019; Long et al., 2018). Given that this has become a mainstream practice in recent fMRI studies, here we seek to reevaluate the original finding with more robust fMRI data sets and to test the generality of this finding to filtered real-world object images as well as artificial object images.

With this rich data set, we found a significant linear correspondence in visual representational structure between the CNNs and the human brain, with a few CNNs (including Alexnet, Cornet-S and Googlenet) exhibiting this relationship across all our image manipulations. This replicated the earlier results (Khaligh-Razavi & Kriegeskorte, 2014; Cichy et al., 2016; see also Güçlü & van Gerven, 2015) and showed that such a brain-CNN relationship is robust even in the presence of various image manipulations and exists for both real-world and artificial object images. This shows that CNNs are able to capture some aspects of neural processing in the human brain.

A linear correspondence between the CNNs and brain, however, only indicates that the representational structure in lower brain regions is better correlated with lower than higher CNN layers and vice versa for higher brain regions, but not how good the correlations are. Although the real-world images we used here were never part of the CNN training set, lower layers of a number of CNNs (including Alexnet, Googlenet, Squeezenet and VGG16) could nevertheless fully capture their representational structure in lower areas of the human visual cortex regardless of whether the original or the filtered versions of these images were shown. CNNs are thus successful in their ability to generalize and compute representations for novel real-world images at lower levels of visual processing in the human brain.

Despite these successes, however, none of the CNNs tested could fully capture the representational structure of the real-world object images at higher levels of visual processing in the human brain. This is in agreement with neurophysiological data showing that the RDM correlation between CNN layers and IT neuronal responses was below the split-half noise ceiling of IT responses (Cadieu et al., 2014; Yamins et al., 2014). When artificial object images were used, not only did most of the CNNs still fail to capture visual processing in higher visual regions, but also none could do so for early visual regions. Thus, despite the presence of a significant linear correspondence in representation between the brain and CNNs, no CNN examined were able to fully capture all levels of visual processing for both real-world and artificial objects. The same results were obtained regardless of whether a CNN was trained with real-world object images or stylized object images that emphasized shape features in its representation.

CNNs implement the known architectures of the primate lower visual processing regions and then repeat this design motif multiple times. The 14 CNNs we examined here were all trained with real-world images from ImageNet (Deng et al., 2009). Their abilities to capture the representations of human lower visual regions when real-world images were used is thus expected. However, the inabilities of such a CNN design motif and training procedure to fully capture the representational structure formed in higher levels of visual processing of real-world objects and both lower and higher levels of visual processing of artificial object images put a significant limit on how CNNs may be used to model the primate vision.

In our fMRI experiments, we used a randomized presentation order for each run of the experiment with two image repetitions. When we simulated the fMRI design precisely in Alexnet by generating a matching number of randomized presentation sequences with image repetitions and then averaging CNN responses for these sequences, we obtained identical Alexnet results as reported here (data not shown). Thus, the disagreement between our fMRI and CNN results could not be due a difference in data processing. The very fact that CNN could fully capture the RDMs in early visual areas during the processing of real-world object images further supports this and additionally shows that the presence of the non-linearities in fMRI measures had a minimal impact on RDM extractions. The latter reflects the robustness of the RSA approach as extensively reviewed elsewhere (Kriegeskorte & Kievit, 2013).

Although we examined object category responses averaged over multiple exemplars rather than responses to each object, previous research has shown similar category and exemplar response profiles in macaque IT and human lateral occipital cortex with more robust responses for categories than individual exemplars due to an increase in SNR (Hung et al., 2005; Cichy et al., 2011). In a recent study, Rajalingham, et al. (2018) found better behavior-CNN correspondence at the category but not at the individual exemplar level. Thus, comparing the representational structure at the category level, rather than at the exemplar level, should have increased our chance of finding a close brain-CNN correspondence. Indeed, the overall brain and CNN correlations for object categories were much higher in the present study than those in previous studies for individual objects (see Khaligh-Razavi & Kriegeskorte, 2014; Cichy et al., 2016). Nevertheless, we found that CNNs failed to fully capture the representational structure of real-world objects in the human brain and performed even worse for artificial objects. Previous studies have shown that object category information is better represented by higher than lower visual regions (e.g., Hong et al., 2016). Our use of object category thus was not optimal for finding a close brain-CNN correspondence at lower levels of visual processing.

Nevertheless, we found better brain-CNN correspondence at lower than higher levels of visual processing for real-world object categories. This suggests that information that defines the different real-world object categories is present at lower levels of visual processing and is captured by both lower visual regions and lower CNN layers. This is not surprising as many categories may be differentiated based on low-level features even with a viewpoint change, such as curvature and the presence of unique features (e.g., the large round outline of a face/head, the protrusion of the limbs in animals) (Rice et al., 2014). Finally, it could be argued that the dissimilarity between the brain and CNNs at higher levels of visual processing for real-world object categories could be driven by feedback from high-level non-visual regions and/or feedback from category-selective regions in human ventral cortex for some of the categories used (i.e., faces, bodies, and houses). However, such feedback should greatly decrease for artificial object categories. Yet we failed to see much improvement in brain-CNN correspondence at higher levels of processing for these objects. If anything, even the strong correlation at lower levels of visual processing for real-world objects no longer existed for these artificial objects.

In recent studies, Baker et al. (2018) and Geirhos et al., (2018, 2019) reported that CNNs rely on local texture and shape features rather than global shape contours. For example, while keeping the contour intact, replacing the skin texture of a cat with that of an elephant resulted in Resnet-50 classifying the animal as an elephant (Geirhos et al., 2018). This may explain in our study why lower CNN layers were able to fully capture the representational structures of real-world object images in lower visual areas, as processing in these brain areas likely relied more on local contours and texture patterns given their smaller receptive field sizes. As high-level object vision relies more on global shape contour processing (Biederman, 1987), the lack of such processing in CNNs may account for CNNs’ inability to fully capture processing in higher visual regions. The artificial objects we used shared similar texture and contour elements at the local level but differed in how these elements were conjoined at the local and global levels. This could explain why none of the CNNs examined could fully capture their representational structure even at the lower level of visual processing in the human brain: presumably a strategy relying on identifying the presence/absence of a particular texture patch is no longer sufficient and human lower visual processing regions likely encode how elements are conjoined at the local level to help differentiate the different objects. Training with stylized images did not appear to improve performance in Resnet-50, suggesting that the differences between CNNs and the human brain may not be overcome by this type of training.

Decades of vision science research has relied on using simple and artificial visual stimuli to uncover the complexity of visual processing in the primate brain, showing that the same algorithms used in processing natural images would manifest themselves in the processing of artificial visual stimuli. The set of artificial object images used in the present study has been used in previous fMRI studies to understand object processing in the human brain (e.g., Op de Beeck et al., 2007; Vaziri-Pashkam & Xu, 2019; Williams et al., 2007). In particular, we recently showed that the transformation of visual representational structures across occipito-temporal and posterior parietal cortices follows a similar pattern for both the real-world objects and the artificial objects used here (Vaziri-Pashkam & Xu, 2019). The disconnection between the representation of real-world and artificial images in CNNs is in disagreement with this long-held principle in primate vision research and suggests that, even during lower levels of visual processing, CNNs may differ from the primate brain in fundamental ways. Such a divergence in lower level processing will undoubtedly contribute to even greater divergence in higher level processing between the primate brain and CNNs.

In our study, we included both shallow and very deep CNNs. Deeper CNNs have been shown to exhibit better object recognition performance (as evident from the ImageNet challenge results, see Russakovsky et al., 2015), and can partially approximate the recurrent processing in ventral visual regions as demonstrated in a recent neurophysiology study (Kar et al., 2019). The recurrent CNN we examined here, Cornet-S, explicitly models recurrent processing that closely correspond to ventral stream visual processing (Kar et al., 2019) and has been argued to be the current best model of the primate ventral visual regions (Kubilius et al., 2018 & 2019). And yet, regardless of these differences, similar performance was observed between shallow and deep CNNs (e.g., Alexnet vs Googlenet), and the recurrent CNN did not perform better than the other CNNs.

Using real-world object images, recent studies have tried to improve brain and CNN RDM correlation by either first finding the best linear transformation between fMRI voxels and CNN layer units and then performing RDM correlations (Khaligh-Razavi et al., 2017) or by using both brain RDM and object categorization to guide CNN training (Kietzmann et al., 2019). Using the first approach, Khaligh-Razavi et al. (2017) reported that the correlation between brain and CNN was able to reach the noise ceiling for LO. A close inspection of the results revealed, however, that brain and CNN correlations were around .2 for all brain regions examined (i.e., V1 to V4 and LO), but noise ceiling decreased from just below .5 in V1 to just below .2 in LO. The low LO noise ceiling again raises concerns about the robustness of this finding (as it did for Khaligh-Razavi & Kriegeskorte, 2014). In Kietzmann et al., (2019), even though brain RDM was incorporated into the training procedure for both a ramping feedforward and a recurrent CNN (with the latter outperforming the former), brain and CNN RDM correlation was still significantly below the noise ceiling in both networks for all the visual regions examined along the human ventral cortex. Thus, the current best model practice (i.e., incorporating brain RDM in CNN training and using a recurrent network architecture) still comes short of fully capturing the representational structure of real-world objects in the human brain. This is likely due to some fundamental differences in visual processing between the brain and CNNs as discussed earlier.

The present results join a host of other studies showing significant differences between the CNNs and brain, such as the kind of features used in object recognition (Ballester & de Araujo, 2016, Ulman et al., 2016; Gatys et al., 2017; Baker et al., 2018; Geirhos et al., 2019;), disagreement in representational structure between CNNs and brain/behavior (Kheradpisheh et al., 2016; Karimi-Rouzbahani et al., 2017; Rajalingham, et al., 2018), the inability of CNN to explain more than 50% of the variance of macaque V4 and IT neurons (Cadieu et al., 2014; Yamins et al., 2014; Kar et al. 2019), and how the two systems handle adversarial images (see Serre, 2019). What the present results add is that even for the key fMRI evidence showing a close brain-CNN correspondence, when more robust data sets and image variation were used, such a correspondence was found to be rather limited and did not fully capture visual processing in the human brain. Most critically, the present results showed that the divergence between the human brain and CNNs increased drastically in how they processed artificial object images. Given that the same algorithms used in processing natural images would manifest themselves in the processing of artificial visual stimuli in the primate brain, this increase in divergence suggests that CNNs likely differ from the primate brain in fundamental ways. CNNs, in their current state, should therefore not be regarded as sound working models of the primate vision in terms of both their explanation power and their potential ability for exploration as proposed by CNN proponents such as Cichy and Kaiser (2019).

Our detailed knowledge of visual processing in the primate early visual areas has contributed significantly to the current success of CNNs. Significant and continuous research effort is critically needed now to uncover the precise algorithms used by the primate brain in visual processing. Doing so could push forward the next leap in model development and make CNNs better models of primate vision. As long as CNNs “see” the world differently than the human brain, they will make mistakes that are against human prediction and intuition. If CNNs are to aid or replace human performance, they need to capture the nature of human vision and then improve upon it. This will ensure both the safety and reliability of the devices powered by CNNs, such as in self-driving cars, and, ultimately, the usefulness of such an information processing system.

## CONCLUSIONS

Presently, the detailed computations performed by CNNs are difficult for humans to understand, rendering them poorly understood information processing systems. By analyzing results from three fMRI experiments and comparing visual representations in the human brain with 14 different CNNs, we found that while CNNs are successful in object recognition, at their current state, they still differ substantially in visual processing from the human brain. This is unlikely to be remedied by an improvement in training, changing the depth of the network, or adding recurrent processing. But rather, some fundamental changes may be needed to make CNNs more brain like. This may only be achieved by our continuous research effort to understand the precise algorithms used by the primate brain in visual processing.

## Materials and Methods

### fMRI Experimental Details

Details of the fMRI experiments have been described in two previously published studies (Vaziri-Pashkam & Xu, 2019 and Vaziri-Pashkam et al., 2019). They are summarized here for the readers’ convenience.

Six, ten, and six healthy human participants with normal or corrected to normal visual acuity, all right-handed, and aged between 18-35 took part in Experiments 1 to 3, respectively. Each main experiment was performed in a separate session lasting between 1.5 and 2 hours. Each participant also completed two additional sessions for topographic mapping and functional localizers. MRI data were collected using a Siemens MAGNETOM Trio, A Tim System 3T scanner, with a 32-channel receiver array head coil. For all the fMRI scans, a T2*-weighted gradient echo pulse sequence with TR of 2 sec and voxel size of 3 mm x 3 mm x 3 mm was used. FMRI data were analyzed using FreeSurfer (surfer.nmr.mgh.harvard.edu), FsFast (Dale et al., 1999) and in-house MATLAB codes. FMRI data preprocessing included 3D motion correction, slice timing correction and linear and quadratic trend removal. Following standard practice, a general linear model was applied to the fMRI data to extract beta weights as response estimates.

In Experiment 1, we used cut-out grey-scaled images from eight real-world object categories (faces, bodies, houses, cats, elephants, cars, chairs, and scissors) and modified them to occupy roughly the same area on the screen (Figure 1B). For each object category, we selected ten exemplar images that varied in identity, pose and viewing angle to minimize the low-level similarities among them. In the original image condition, unaltered images were shown. In the controlled image condition, images were shown with contrast, luminance and spatial frequency equalized across all the categories using the SHINE toolbox (Willenbockel et al., 2010, see Figure 1B). Participants fixated at a central red dot throughout the experiment. Eye-movements were monitored in all the fMRI experiments to ensure proper fixation.

During the experiment, blocks of images were shown. Each block contained a random sequential presentation of ten exemplars from the same object category. Each image was presented for 200 msec followed by a 600 msec blank interval between the images (Figure 1A). Participants detected a one-back repetition of the exact same image. This task engaged participants’ attention on the object shapes and ensured robust fMRI responses. However similar visual representations may be obtained when participants attended to the color of the objects (Vaziri-Pashkam & Xu, 2017, Xu & Vaziri-Pashkam, 2019; see also Xu, 2018). Two image repetitions occurred randomly in each image block. Each experimental run contained 16 blocks, one for each of the 8 categories in each image condition (original or controlled). The order of the eight object categories and the two image conditions were counterbalanced across runs and participants. Each block lasted 8 secs and followed by an 8-sec fixation period. There was an additional 8-sec fixation period at the beginning of the run. Each participant completed one scan session with 16 runs for this experiment, each lasting 4 mins 24 secs.

In Experiment 2, only six of the original eight object categories were included and they were faces, bodies, houses, elephants, cars, and chairs. Images were shown in 3 conditions: Full-SF, High-SF, and Low-SF. In the Full-SF condition, the full spectrum images were shown without modification of the SF content. In the High-SF condition, images were high-pass filtered using an FIR filter with a cutoff frequency of 4.40 cycles per degree (Figure 1B). In the Low-SF condition, the images were low-pass filtered using an FIR filter with a cutoff frequency of 0.62 cycles per degree (Figure 1B). The DC component was restored after filtering so that the image backgrounds were equal in luminance. Each run contained 18 blocks, one for each of the category and SF condition combination. Each participant completed a single scan session containing 18 experimental runs, each lasting 5 minutes. Other details of the experiment design were identical to that Experiment 1.

In Experiment 3, we used unaltered images from both real-world and artificial object categories. The real-world categories were the same eight categories used in Experiment 1. The artificial object categories were nine categories of computer-generated 3D shapes (ten images per category) adopted from Op de Beeck et al. (2008) and shown in random orientations to increase image variation within a category (see Figure 5A). Each run of the experiment contained 17 stimulus blocks, one for each object category (either real-world or artificial). Each participant completed 18 runs, each lasting 4 mins 40 secs. Other details of the experiment design were identical to that Experiment 1.

We examined responses from independent localized early visual areas V1 to V4 and higher visual processing regions LOT and VOT. V1 to V4 were mapped with flashing checkerboards using standard techniques (Sereno et al., 1995). Following the detailed procedures described in Swisher et al. (2007) and by examining phase reversals in the polar angle maps, we identified areas V1 to V4 in the occipital cortex of each participant (see also Bettencourt & Xu, 2016) (Figure 1C). To identify LOT and VOT, following Kourtzi and Kanwisher (2000), participants viewed blocks of face, scene, object and scrambled object images. These two regions were then defined as a cluster of continuous voxels in the lateral and ventral occipital cortex, respectively, that responded more to the original than to the scrambled object images (Figure 1C). LOT and VOT loosely correspond to the location of LO and pFs (Malach et al., 1995; Grill-Spector et al.,1998; Kourtzi & Kanwisher, 2000) but extend further into the temporal cortex in an effort to include as many object-selective voxels as possible in occipito-temporal regions.

To generate fMRI response patterns for each object category for the main experiments in each condition and in each run, we first convolved an 8-second stimulus presentation boxcar (corresponding to the length of each image block) with a hemodynamic response function. We then conducted a general linear model analysis to extract beta value for each condition in each voxel of each ROI. This was done separately for each run. We normalized the beta values across all voxels of a given ROI in a given run using z-score transformation to remove amplitude differences between runs, conditions and ROIs. Following Tahan and Konkle (2019), we selected the top 75 most reliable voxels in each ROI. This was done by splitting the data into odd and even halves, correlating all the conditions in an experiment between the two halves for each voxel, and selecting the top 75 voxels showing the highest correlation. This is akin to including the best units in monkey neurophysiological studies. For example, Cadieu et al. (2014) only selected a small subset of all recorded single units for their brain-CNN analysis.

### CNN details

We included 14 CNNs in our analyses (see Table 1). They included both shallower networks, such as Alexnet, VGG16 and VGG 19, and deeper networks, such as Googlenet, Inception-v3, Resnet-50 and Resnet-101. We also included a recurrent network, Cornet-S, that has been shown to capture the recurrent processing in macaque IT cortex with a shallower structure (Kubilius et al., 2019; Kar et al., 2019). This CNN has been recently argued to be the current best model of the primate ventral visual processing regions (Kar et al., 2019). All the CNNs used were trained with ImageNet images (Deng et al., 2009).

To understand how the specific training images would impact CNN representations, besides CNNs trained with ImageNet images, we also examined Resnet-50 trained with stylized ImageNet images (Geirhos et al., 2019). We examined the representations formed in Resnet-50 pretrained with three different procedures (Geirhos et al., 2019): trained only with the stylized ImageNet Images (RN50-SIN), trained with both the original and the stylized ImageNet Images (RN50-SININ), and trained with both sets of images and then fine-tuned with the stylized ImageNet images (RN50-SININ-IN).

Following O’Connor et al. (2018), we sampled between 6 and 11 mostly pooling and FC layers of each CNN (see Table 1 for the specific CNN layers sampled). Pooling layers were selected because they typically mark the end of processing for a block of layers before information is pooled and passed on to the next block of layers. When there were no obvious pooling layers present, the last layer of a block was chosen. For a given CNN layer, we extracted the CNN layer output for each object image and then averaged the output from all images in a given category to generate the CNN layer response for that object category. Cornet-S and the different versions of Resnet-50 were implemented in Python. All other CNNs were implemented in Matlab. Output from all CNNs were analyzed and compared with brain responses using Matlab.

### Comparing the representational structures between the brain and CNNs

To determine the extent to which object category representations were similar between brain regions and CNN layers, we correlated the object category representational structure between brain regions and CNN layers. To do so, we obtained the representational dissimilarity matrix (RDM) from each brain region by computing all pairwise Euclidean distances for the object categories included in an experiment and then taking the off-diagonal values of this RDM as the category dissimilarity vector for that brain region. This was done separately for each participant. Likewise, from the CNN layer output, we computed pairwise Euclidean distances for the object categories included in an experiment to form the RDM and then taking the off-diagonal values of this RDM as the category dissimilarity vector for that CNN layer. We applied this procedure to each sampled layer of each CNN. We then correlated the category dissimilarity vectors between each brain region of each participant and each sampled CNN layer. Following Cichy et al. (2016), all correlations were calculated using Spearman rank correlation to compare the rank order, rather than the absolute magnitude, of the category representational similarity between the brain and CNNs (see also Nili et al., 2014). All correlation coefficients were Fisher z-transformed before group-level statistical analyses were carried out.

To evaluate the correspondence in representation between lower and higher CNN layers to lower and higher visual processing regions, for each CNN examined, we identified in each human participant, the CNN layer that showed the best RDM correlation with each of the six brain regions included. We then assessed whether the resulting layer numbers linearly increased from low to high visual regions using Spearman rank correlation. Finally, we tested the resulting correlation coefficients at the group level. If a close correspondence in representation exists between the brain and CNNs, the averaged correlation coefficients should be significantly above zero. All stats reported were corrected for multiple comparisons for the number of comparisons included in each experiment using the Benjamini–Hochberg procedure with false discovery rate (FDR) controlled at q = 0.05 (Benjamini & Hochberg, 1995).

To assess how successfully the category RDM from a CNN layer could capture the RDM from a brain region, we first obtained the reliability of the category RDM in a brain region across the group of human participants by calculating the lower and upper bounds of the noise ceiling of the fMRI data following the procedure described by Nili et al. (2014). Specifically, the upper bound of the noise ceiling for a brain region was established by taking the average of the correlations between each participant’s RDM and the group average RDM including all participants, whereas the lower bound of the noise ceiling for a brain region was established by taking the average of the correlations between each participant’s RDM and the group average RDM excluding that participant.

To evaluate the degree to which CNN category RDMs may capture those of the different brain regions, for each CNN, using t tests, we examined how close the highest correlation between a CNN layer and a brain region was to the lower bound of the noise ceiling of that brain region. The t test results were corrected for multiple comparisons for the 6 brain regions included using the Benjamini–Hochberg procedure. If a CNN layer was able to fully capture the representational structure of a brain region, then its RDM correlation with the brain region should exceed the lower bound of the noise ceiling of that brain region. Because the lower bound of the noise ceiling varied somewhat among the different brain regions, for illustration purposes, we plotted the lower bound of the noise ceiling from all brain regions at 0.7 while maintaining the differences between the CNN and brain correlations with respect to their lower bound noise ceilings. This did not affect any statistical test results.

## Acknowledgement

We thank Martin Schrimpf for help implementing CORnet-S, JohnMark Tayler for extracting the features from the three Resnet-50 models trained with the stylized images, and Thomas O’Connell, Brian Scholl, JohnMark Taylor and Nick Turk-Brown for helpful discussions and feedback on the results. The fMRI data collected was supported by NIH grant 1R01EY022355 to YX.

## Supplemental Figure Captions

**Supplemental Figure 1.**
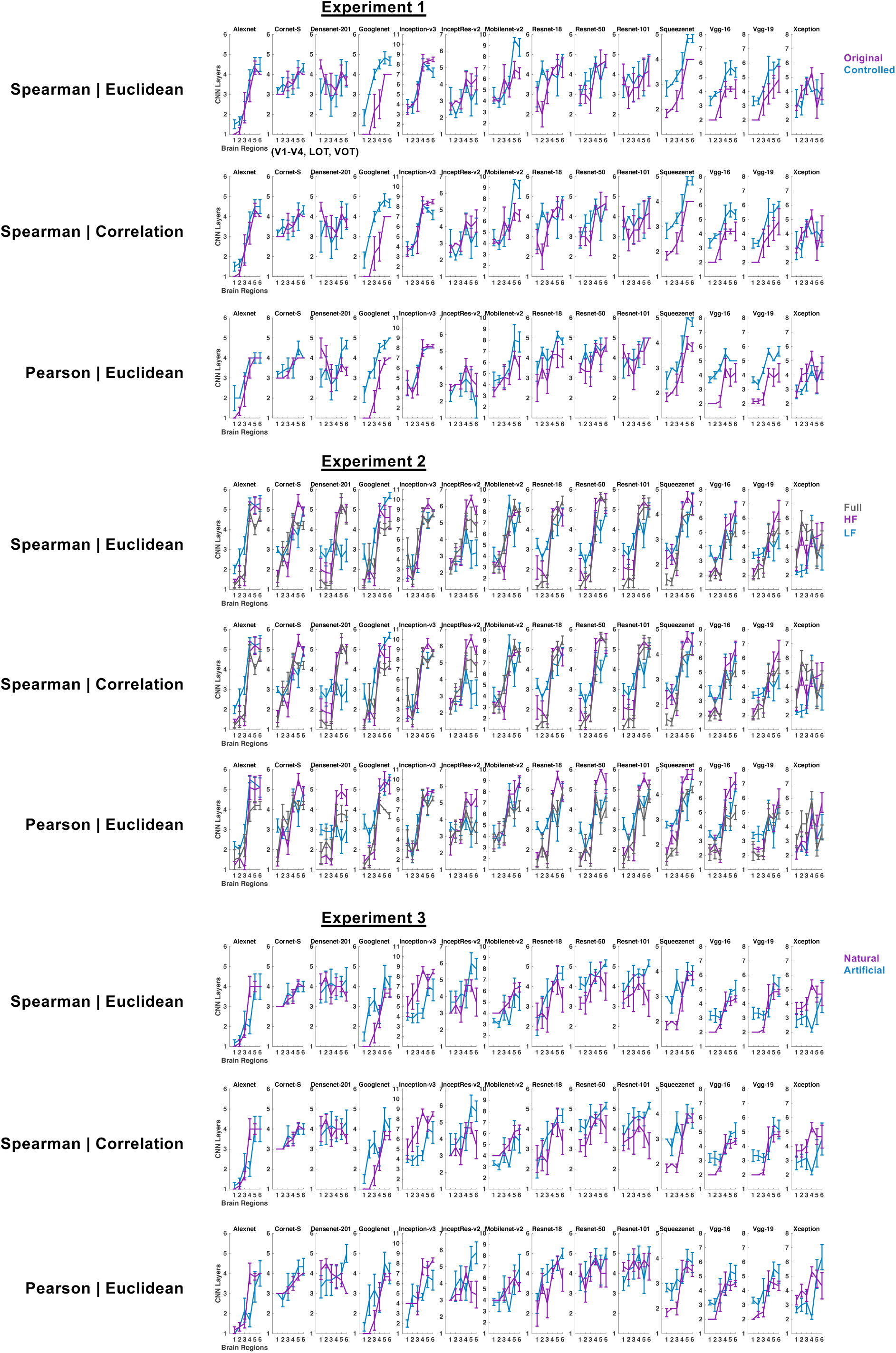
Evaluating the presence of the brain-CNN correspondence in their representational structures for Experiments 1 to 3 using three different measures: Spearman correlation with Euclidean distance measures (same as those shown in Figure 2), Spearman correlation with correlation measure; and Pearson correlation with Euclidean distance measures. Virtually the same results were obtained in all three measures.

**Supplemental Figure 2.**
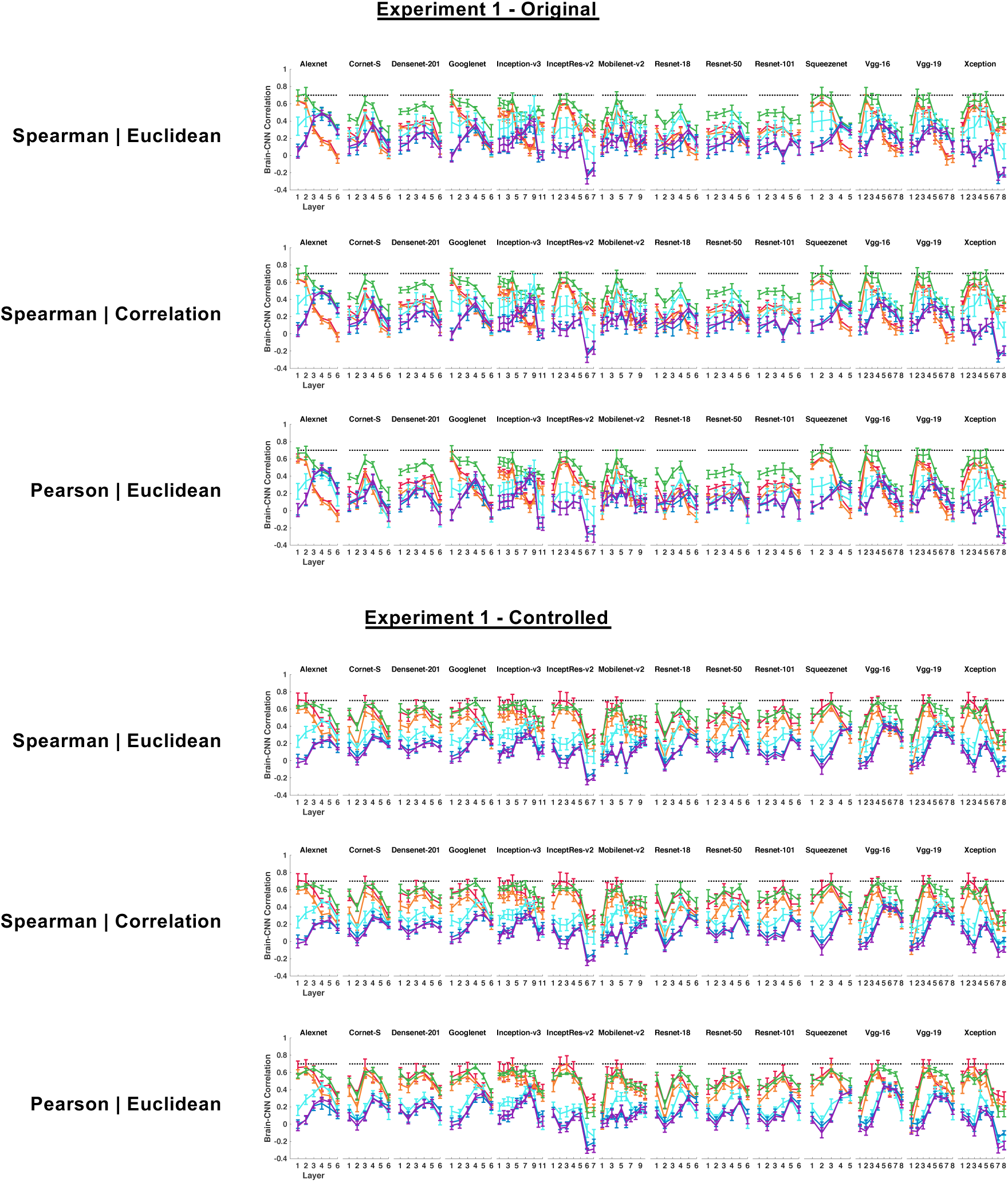
Quantifying the brain-CNN correspondence in Experiment 1 using three different measures: Spearman correlation with Euclidean distance measures (same as those shown in Figure 2), Spearman correlation with correlation measure; and Pearson correlation with Euclidean distance measures. Virtually the same results were obtained in all three measures.

**Supplemental Figure 3.**
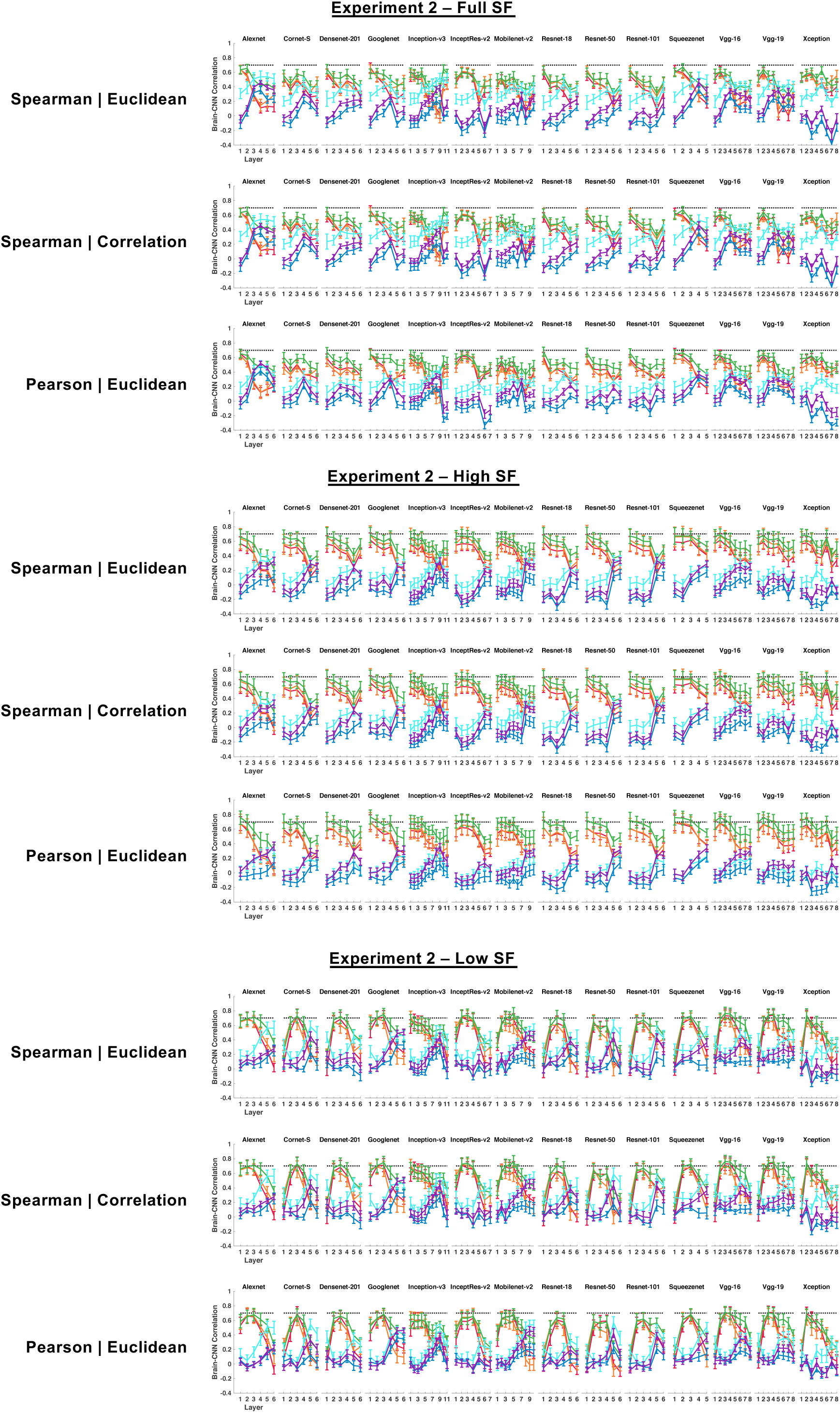
Quantifying the brain-CNN correspondence in Experiment 2 using three different measures: Spearman correlation with Euclidean distance measures (same as those shown in Figure 2), Spearman correlation with correlation measure; and Pearson correlation with Euclidean distance measures. Virtually the same results were obtained in all three measures.

**Supplemental Figure 4.**
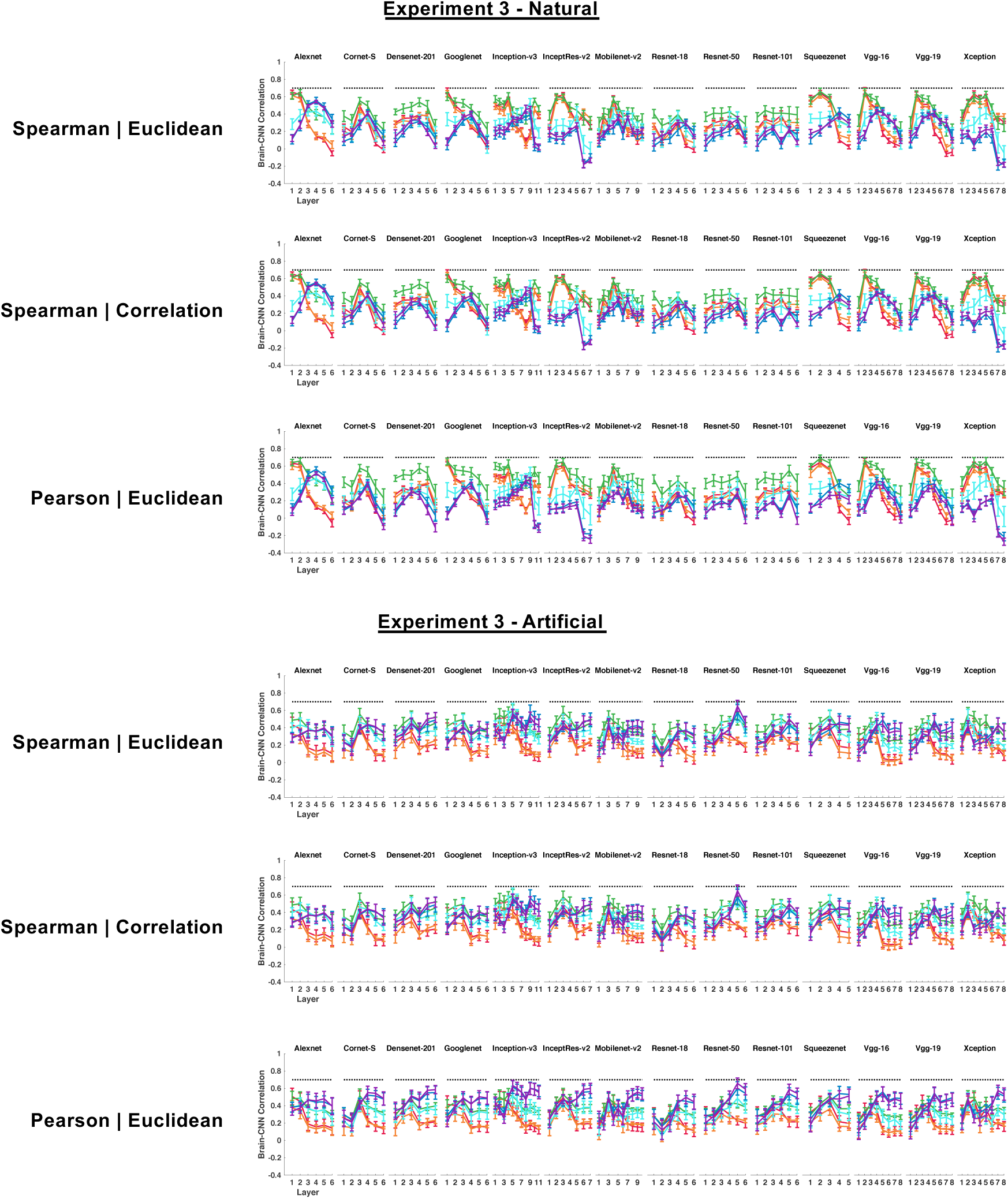
Quantifying the brain-CNN correspondence in Experiment 3 using three different measures: Spearman correlation with Euclidean distance measures (same as those shown in Figure 2), Spearman correlation with correlation measure; and Pearson correlation with Euclidean distance measures. Virtually the same results were obtained in all three measures.

**Supplemental Figure 5.**
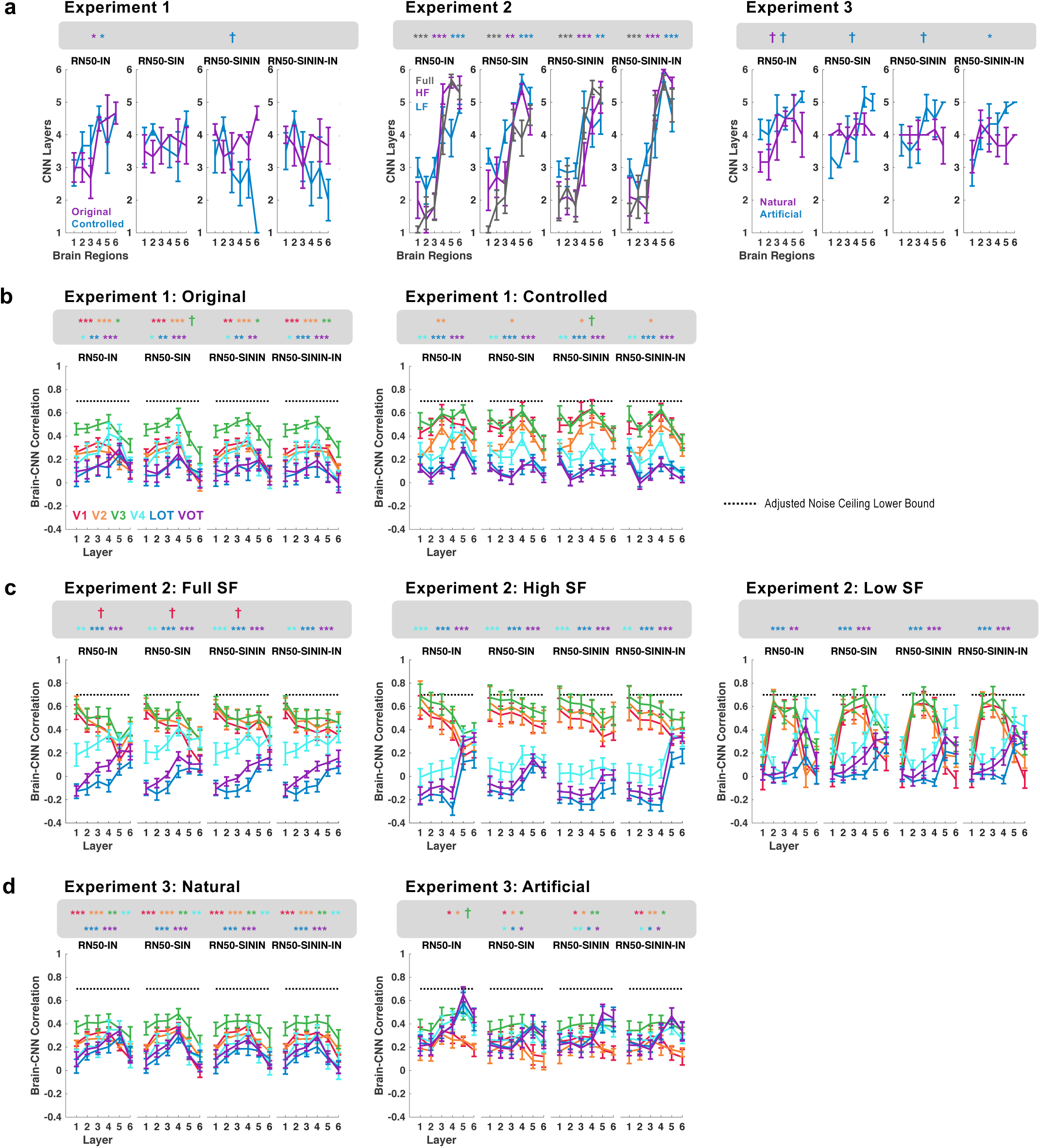
Comparing Resnet-50 trained with original and stylized ImageNet images. (A) The correspondence between the brain and CNN in their representational structure. (B) to (D) RDM correlations of each brain region with each sampled layer in each CNN for the images in Experiments 1 to 3, respectively. Resnet-50 was pretrained either with the original ImageNet images (RN50-IN), the stylized ImageNet Images (RN50-SIN), both the original and the stylized ImageNet Images (RN50-SININ), or both sets of images and then fine-tuned with the stylized ImageNet images (RN50-SININ-IN). The lower bounds of the noise ceiling from all brain regions were set to 0.7. † p < .1, * *p < .05*, ** *p < .01*, *\*\*\* p < .001*.

